# The Rab32 small GTPase is required for efficient cross-priming of CD8^+^ T cells against cell-associated antigens by XCR1^+^ type 1 DCs *in vivo*

**DOI:** 10.1101/2025.05.03.652057

**Authors:** Naman Tandon, Enric Gutiérrez-Martínez, Pierre Bourdely, Giorgio Anselmi, Jordan Denisot, Yohan Gerber, Julie Helft, Marcelle Bens, Loredana Saveanu, Pierre Guermonprez

## Abstract

Type 1 conventional dendritic cells (cDC1s) are critical for initiating adaptive immune responses through the cross-priming of CD8⁺ T cells against antigens from tumor or virus-infected cells.

This function depends on specialized cross-presentation pathways that allow cDC1s to process phagocytosed cell debris and present peptide–MHC I complexes. In this study, we identify the small GTPase Rab32 as being highly and selectively over-expressed in cDC1s as compared to cDC2s. While cDC1s from Rab32-deficient mice develop normally and can respond to maturation signals, their capacity to activate CD8⁺ T cells *in vivo* is impaired. Notably, Rab32- deficient cDC1s retain the ability to stimulate TCR transgenic CD8⁺ T cells ex vivo using both cell-associated antigens and MHC I-binding peptides of varying affinity.

However, *in vivo*, Rab32 is essential for effective CD8⁺ T cell responses to cell-associated antigens, independent of Rab32 expression in T cells themselves. Importantly, Rab32-mediated cross-priming is required for the efficient expansion of tumor-specific CD8⁺ T cells into solid tumors. These findings underscore a critical role for Rab32 in cDC1-mediated cross-priming, highlighting the contribution of non-antigen processing vesicular pathways in shaping CD8⁺ T cell responses to cellular antigens.

## Introduction

Conventional dendritic cells (cDCs) constantly survey lymphoid as well as peripheral tissues in the body for antigen-expressing cells such as necrotic cells, pathogen-infected or tumor cells. Upon such encounters, cDCs internalize and process the cell-associated protein antigens, and present them as peptides on MHC-I to naïve CD8^+^ T cells in a process termed as cross-presentation (1)(2). When accompanied by co-stimulatory and cytokine signals (IL-12, type-1 IFNs, IL-15, *e.g.*), cross-presentation leads to the differentiation of antigen-specific CD8^+^ T cells into effector T cells.

cDCs are further divided into two subsets in mice as well as humans – XCR1^+^ cDC1s and SIRPa^+^ cDC2s. In mice, cDC1s are functionally specialized at cross-presentation of cell-associated antigens and are critical for cross-priming against viral and tumoral antigens *in vivo* (3)(4)(5)(6)(7). This specialization seems to be conserved in humans since human blood-derived as well as dermal human cDC1s can also efficiently cross-present cell-associated antigens *in vitro* (8–11).

Several properties of cDC1s support their specialization at cross-presentation over cDC2s. cDC1s seem to be more efficient at internalizing necrotic cell-associated antigens compared to cDC2s *in vivo* (12). Similarly, migratory cDC1s in the lung exclusively pick up necrotic cell-associated antigens (13)as well as viral antigens from infected cells (5) and transport them to draining lymph nodes for cross-presentation *in vivo*. Independently of antigen uptake, Schnorrer et al. have shown that cDC1s possess a superior ability to cross-present phagocytosed antigens as compared to cDC2s (7).

While cDC1s are the primary cross-presenting cells in physiological settings, a significant fraction of studies investigating the cellular mechanism of cross-presentation have been conducted in murine bone marrow-derived DCs (BMDCs), generated *in vitro* from GM-CSF supplemented bone marrow cultures (14)(15)(16)(17). BMDCs are a heterogeneous population containing both DCs and macrophages, and they use a cross-presentation transcriptional program different from cDC1s (18).(19). Hence, multiple studies have investigated the cross-presentation machinery utilized by cDC1s. Several cDC1 enriched proteins have been implicated in facilitating cross- priming of CD8^+^ T cells against cellular antigens *in vivo*, including Clec9a (20)(21) Perilipin2 (22), Rab43 (23), WDFY4 (24), Rab39a (25), Perforin-2 (26) and APOL7C (27).

It is believed that special features of the endocytic pathway of cDC1 might promote the conservation of internalized antigens in the phagosomes, and their efficient delivery in the cytosol (28)(26) (27) thus allowing optimal processing of antigens into MHC-I compatible peptides. Other processes, such as Perilipin 2-dependent accumulation of neutral lipids (22) and expression of WDFY4 (24) contribute to cross-presentation by mechanisms that are poorly understood. Finally, cDC1s show enhanced expression of the MHC-I peptide loading complex, which supports their superior cross-presentation abilities (29).

Rab proteins are small GTPases that are well conserved in mammals, and they make interesting candidates for cross-presentation studies in cDC1s. These proteins function as the master regulators of intracellular vesicular trafficking. On these lines, Rab GTPases have been shown to facilitate cross-presentation by regulating the trafficking events involved in antigen processing and peptide: MHC-I loading (30)(15,23,25,31). The GTPase Rab32 plays a critical role in the restriction and clearance of intracellular bacterial pathogens such as *Listeria monocytogenes* (*32*) and *Salmonella* (33)(34) (35) within DCs and macrophages, respectively. The itaconate metabolism is downstream Rab32 (36). Rab32 deficiency in CD11c^+^ cells has also been linked to increased colitis progression and bacterial invasion in colon tissues (37). Intracellular analysis in cell lines shows that Rab32 colocalizes with the mitochondria-associated membranes of the ER (MAMs), which are marked by the ER chaperone calnexin (38)(39)(40). Moreover, Rab32 and its guanine-nucleotide exchange factor (GEF) BLOC-3 (HPS1-4) play an essential role in the biogenesis of lysosome-related organelles (LROs) (41)(42). Disruptions in this machinery have been linked to Hermansky-Pudlak syndrome (HPS) in humans, which is characterized by oculocutaneous albinism, bleeding tendency, and immunodeficiencies. The role of Rab32 in adaptive immunity has not yet been critically investigated.

In this report, we show that Rab32 is highly expressed in cross-presenting cDC1s and investigate its role in the cross-priming of CD8^+^ T cells in response to cross-presentation of cell-associated antigens by cDC1s.

## Materials and methods

*Mice* – All mice experimentation was performed under authorisation from the Ministere de l’education nationale, de l‘enseignement superieur et de la recherche, APAFIS n. 15373 - 2018100811239028 v6. The Rab32 tm1a mice have a promoter-driven cassette inserted between exon 1 and 2 of Rab32, which disrupts Rab32 translation in these mice. They have a CD45.2^+^ C57BL/6 background and were purchased from the Knockout Mice Project (KOMP), UC Davis, US. To generate littermates, Rab32 tm1a mice were crossed with WT C57BL/6 mice purchased from Charles River, France. WT/WT and tm1a/tm1a littermates were identified by genotyping in the second progeny generation and were used for experimentation. The Rab32 cdel mice have a constitutive deletion of Rab32 exon 2. They were generated at CNRS TAAMS at Orleans, France, by crossing the CD45.2^+^ Rab32 tm1a mice (possessing floxed exon2) with Cd45.2^+^ germ-line cre expressing ‘Rosa26 Deleter CRE’ mice. WT/WT and cdel/cdel littermates were identified by genotyping in the second progeny generation and were used for experimentation. The CD45.1.2^+^ OT-I mice were generated in-house by crossing male CD45.2^+^ OT-I Rag2 knockout mice (a kind gift from Dr Oliver Lantz, Institut Curie, France) with female CD45.1^+^ OT-II mice (purchased from Charles River Laboratories). The first generation CD45.1.2^+^ OT-I mice were then used for experimentation. The genotyping primers for Rab32 tm1a mice (cassette region) are – a. TGCAGGCAGTAGGCATTCTA, b. CCAACTGACCTTGGGCAAGAACAT, The genotyping primers for Rab32 cdel mice (exon 2 deletion) are – a. CACACCTCCCCCTGAACCTGAAA, b. TGTTTTCTTGGCCTCTTTCAA. The genotyping primers for the cre gene are – a. CCTGGAAAATGCTTCTGTCCG, b. CAGGGTGTTATAAGCAATCCC.

*Cell lines* – K^bm1^/K^bm1^_OVA MEFs were a kind gift from Dr Caetano Reis e Sousa, The Francis Crick Institute, London. These cells were cultured in complete DMEM 1X medium (Gibco™ Life Technologies), supplemented with 10% fetal bovine serum (FBS) (Gibco™ Life Technologies), 1% penicillin-streptomycin (Gibco™ Life Technologies) and 0.1 % 2-Mercaptoethanol (Sigma-Aldrich). The B16/ B16_OVA/ B16-FLT3L melanomas were cultured in a similar preparation in RPMI medium (Gibco™ Life Technologies). The cells were passaged every 2-3 days depending on the confluence using the 2.5% 10X Trypsin solution (Gibco™ Life Technologies).

*Flow Cytometry*- Flow cytometry was performed on the BD LSRFortessa^TM^ FACS instrument. All data was analysed using FlowJo analysis software (Tree Star). Antibody staining was performed at on ice in home-made FACS buffer (PBS + 2% BSA + 2mM EDTA). The list of antibodies used for flow cytometry analysis are provided in supplementary materials.

*Purification and sorting of primary cDC1s* – Mice were injected with B16-FLT3L to promote the numbers of cDC1s recovered. On day 9 after tumor injection, spleens were harvested from the mice and digested in a 0.375 U/ ml solution of Collagenase D (ROCHE Ref 11-088-866-001, Lot n. 20350022) in the digestion medium (HBSS, 5 % FCS, b-me 50uM) for 30 min at 37^0^C. RBCs were removed by treatment with home-made ACK lysis buffer (NH4Cl= 8290mg/L, KHCO_3_= 5000mg/L, EDTA= 0.0995mM in 1L distilled water). Splenocytes were then passed through a 70um cell strainer, pelleted and stained with a home-made antibody cocktail for 30 min at 4^0^C (list of antibodies provided in supplementary materials). Anti-biotin microbeads (Miltenyi Biotec 130- 090-485) were then added (10ul per 10^7^ cells as per manufacturer’s protocol), and the solution was incubated for 10 min at 4^0^C. The solution was then passed through an LS MACS column (Miltenyi Biotec 130-042-401) placed in a magnetic field, and the flow-through enriched in XCR1^+^ CD11c^+^ cDC1s was collected. The XCR1^+^ CD11c^+^ cDC1s were further sorted using the BD FACSMelody™ Cell Sorter. Sorted cells were resuspended in complete IMDM (supplemented with 10% FBS, 1% penicillin-streptomycin and 0.1 % 2-Mercaptoethanol).

*Preparation of OT-I cells* – Various lymph nodes were collected from OT-I mice in a 6 well plate on ice, including the inguinal, axillary, brachial and mesenteric lymph nodes. They were teased open to bring out the lymphocytes, and then gently smashed onto a 70um cell strainer within the well. The well and the cell strainer were properly washed with cold FACS buffer, and lymphocyte solution was were pelleted by centrifugation. The cells were labelled with 5uM CTV (Life technologies C34557) in PBS 0.1% BSA. The OT-I cells were then isolated using the CD8a^+^ T cell isolation kit (Miltenyi Biotec 130-104-075) by following the manufacturer’s protocol. For injections, the cells were responded in sterile PBS, whereas for culture the cells were resuspended in complete IMDM medium (supplemented with 10% FBS, 1% penicillin-streptomycin and 0.1 % 2-Mercaptoethanol).

*Poly(I:C) preparation for injections-* For annealing the poly(I:C) HMW (Invivogen), required concentration was heated at 65°C in water bath for 10 min, then allowed to cool down at RT for 45 min. Annealed poly(I:C) was then used as required.

*Assessment of endogenous CD8^+^ T cells*– K^bm1^/K^bm1^_OVA cells were passaged and a cell solution was prepared in PBS at 1x10^6^ cells/180ul/injection. The cells were UV irradiated at 9999uJoules x 100 for 10 minutes. 10ug/20ul/injection annealed poly(I:C) was added to the cell solution. 200ul solution (K^bm1^/K^bm1^_OVA+10ug poly(I:C)) was *i.v.* injected in *Rab32^+/+^* or *Rab32*^-/-^ mice. 10 days later, 500ul blood was collected from mice by cardiac puncture in the presence of heparin (anti- coagulant), following which the mice were sacrificed and their spleens were harvested. Both blood and spleen samples were treated with ACK buffer to lyse RBCs, counted by flow cytometry using Accucheck counting beads (Life Technologies PCB100) and stained with iTAg Tetramer/PE - H- 2^Kb^ OVA (SIINFEKL) (1:20 dilution, MBL TB-5001-1) for 30 min at RT. Antibody mix was then added and the cells were stained for 30 min on ice (list of antibodies provided in supplementary

materials). The samples were washed twice in cold PBS by centrifugation, and stained with Streptavidin BV605 for lineage negative cells (1:500 dilution, BD Biosciences 563260) for 30 min on ice. The cells were then washed twice in PBS by centrifugation, analysis by flow cytometry.

*OT-I adoptive transfer experiment* – CD45.1.2^+^ OT-I cells were CTV labelled and purified as described above. 1x10^6^ cells were *i.v.* injected in CD45.2^+^ host *Rab32^+/+^* or *Rab32*^-/-^ mice. 6-12h later, different numbers of live K^bm1^_OVA fibroblasts were *i.v.* injected in the same mice. On day 3 p.i., mice were sacrificed and their spleens were harvested, treated with ACK buffer, counted and stained for flow cytometry analysis.

*Ex vivo peptide presentation assay –* 10,000 splenic XCR1^+^CD11c^+^ cDC1s were counted and plated per condition. The SIINFEKL and SIIQFEKL peptides (JPT peptide technology, Germany) were serially diluted in complete IMDM (supplemented with 10% FBS, 1% penicillin-streptomycin and 0.1 % 2-Mercaptoethanol), and different concentrations were added to the respective wells. 100,000 purified and CTV labelled OT-I cells were then added to all the wells. The plates were incubated at 37^0^C, and 24h later the culture supernatant was collected and stored in -20^0^C for IL-2 ELISA, and the culture medium was replenished in the wells. 48h later (day 3 of culture), the supernatant was collected for IFNγ ELISA. The cells were stained and analysed by flow cytometry.

*Ex vivo cross-presentation assay* – XCR1^+^CD11c^+^ cDC1s were sorted from the spleens of *Rab32^+/+^* or *Rab32*^-/-^ mice as described above. 10,000 cells were counted and plated per condition. Different numbers of K^bm1^_OVA cells were added to the respective cells. 100,000 purified OT-I cells were then added to all the wells. The plates were incubated at 37^0^C for 24h, and culture supernatant was collected and stored in -20^0^C. This supernatant was used to perform IL-2 and IFNγ ELISA using the manufacturer’s protocol (Mouse IFNγ ELISA MAX^TM^ Standard Set, Biolegend 430801 and Mouse IL-2 ELISA MAX^TM^ Deluxe set, Biolegend 431004).

*Tumor OT-I experiment-* CD45.2^+^ host *Rab32^+/+^* or *Rab32*^-/-^ mice were *s.c.* injected with 0.5x10^6^ B16-OVA cells. The tumors were allowed to grow to measurable sizes, and on day11-14, 1x10^6^ purified CD45.1.2^+^ OT-I cells were injected i.v. in the same mice. On day 17-16, the mice were sacrificed, and the tumors were harvested. Tumors were then digested with Collagenase D (ROCHE Ref 11-088-866-001) in the digestion medium (HBSS, 5 % FCS, b-me 50uM) for 40 min at 37^0^C, and RBC lysis was performed as described above. 5x10^6^ CD45^+^ cells were then counted using Accucheck beads by flow cytometry, plated and stained for further flow cytometry analysis.

*Confocal microscopy –* Glass coverslips were coated with purified I-A/I-E antibody (100 dil, Biolegend 107601) overnight at 4^0^C. The next day, 30,000 sorted splenic XCR1^+^ CD11c^+^ cDC1s were added per coverslip and were allowed to spread for 2h at 37^0^C. The cells were fixed in 2% paraformaldehyde (PFA) for 15 min at 37^0^C, followed by two washes in PBS. The cells were then treated with the permeabilisation solution (PBS+0.2% Saponin+0.2% BSA) at RT for 20 min. Primary antibody staining was performed in the permeabilisation solution at RT for 40 min. The cells were then washed twice in PBS, and stained for secondary antibodies along with 1X DAPI for nuclear staining, for 40 min at RT. Antibody references are provided in supplementary information. Following two washes, a final fixation was performed in 4% formaldehyde in PBS. The samples were then treated with 50mM NH_4_Cl for 7 min at RT for quenching the residual formaldehyde, washed and mounted on glass slides with fluoromount mounting medium. Images were acquired on Leica Sp8 confocal microscope (Leica Microsystems, Germany) using the 63x immersion lens, and deconvoluted using the Huygens Professional image processing software. Further processing was performed using the Fiji-ImageJ software.

*Western blot –* Approximately 500,000 cells were washed in PBS and pelleted by centrifugation for analysis. The pellet was resuspended in 40ul of cell lysis buffer comprising 0.5% NP40 and 1X protease inhibitor (Roche 05892988001) in PBS. Cell lysis was performed for 30 min on ice, accompanied by vortexing after every 10 min. The solution was centrifuged for 20 min at 15000 rpm at 4°C, following which the supernatant was carefully collected in a separate tube. Cell extract was denatured in Laemmli buffer at 95^0^C for 5 min and then transferred on ice for 5 min. The tube was centrifuged at 2000rpm for 2 min, and the supernatant containing the denatured cell extract was loaded on precast polyacrylamide gels (4–15% Bio-Rad Criteron TGX Stain-Free gels, Cat n. 5678084), along with the molecular ladder. After running the gel, the blots were transferred to a nitrocellulose membrane. Non-specific reactions were blocked by incubating the membrane with Tris-buffered solution (TBS) containing 0.5% Tween-20 (TBS.T) and 5% milk at 4^0^C overnight. The following day, the membrane was cut into two parts just below the 37KDa mark to perform separate stainings for Rab32 (25KDa) and Actin (42KDa). The membranes were stained with respective primary antibodies in TBS.T for 1.5h at RT - Rab32 (400 dil, host mouse, Santa Cruz 390178) or Actin (20,000 dil, host mouse, Sigma A1978). Following 3 washes in TBS.T with 10 min incubations each, the membranes were stained with secondary anti-Mouse HRP conjugated IgG (H+L) (10000 dil, Thermofisher 62-6520) for 1h on the shaker at RT. Following 3 washes in TBS.T, the membranes were developed with the Clarity Western ECL Substrate (Bio-Rad 1705061) and analysed within 15 min using Bio-Rad ChemiDoc™ Imager. The blot images were processed in the ImageLab software.

*Gene microarray data* – The data was downloaded from the Immunological genome project database at Immgen.org. Heat maps were created using the web-based Morpheus tool (Broad Institute - https://software.broadinstitute.org/morpheus).

*Statistical analysis* – All statistical analyses were performed using the Prizm GraphPad software (Graphpad Software Inc., San Diego, CA, USA).

## Results

### Rab32 is highly expressed in cDC1s as compared to cDC2s or pDCs but does not control their differentiation

To define the potential regulators of cross-priming, we sought to identify cDC1-enriched Rab GTPases by evaluating the expression profile of Rab proteins in different populations of murine DC subsets. To this end, we looked at the gene expression microarray data from the Immgen database. As compared to cDC2s or pDCs, cDC1s highly expressed Rab32 (Fig 1a). This was observed in spleen, mesenteric (MLN) and skin draining lymph node (SLN) (Fig 1a). In addition, Rab32 is also expressed in some other myeloid populations (alveolar macrophages, *e.g*.) but not in lymphocytes in mice (Fig S1a). In conclusion, Rab32 is mainly expressed in cDC1s amongst professional antigen-presenting cells.

**Figure 1.**
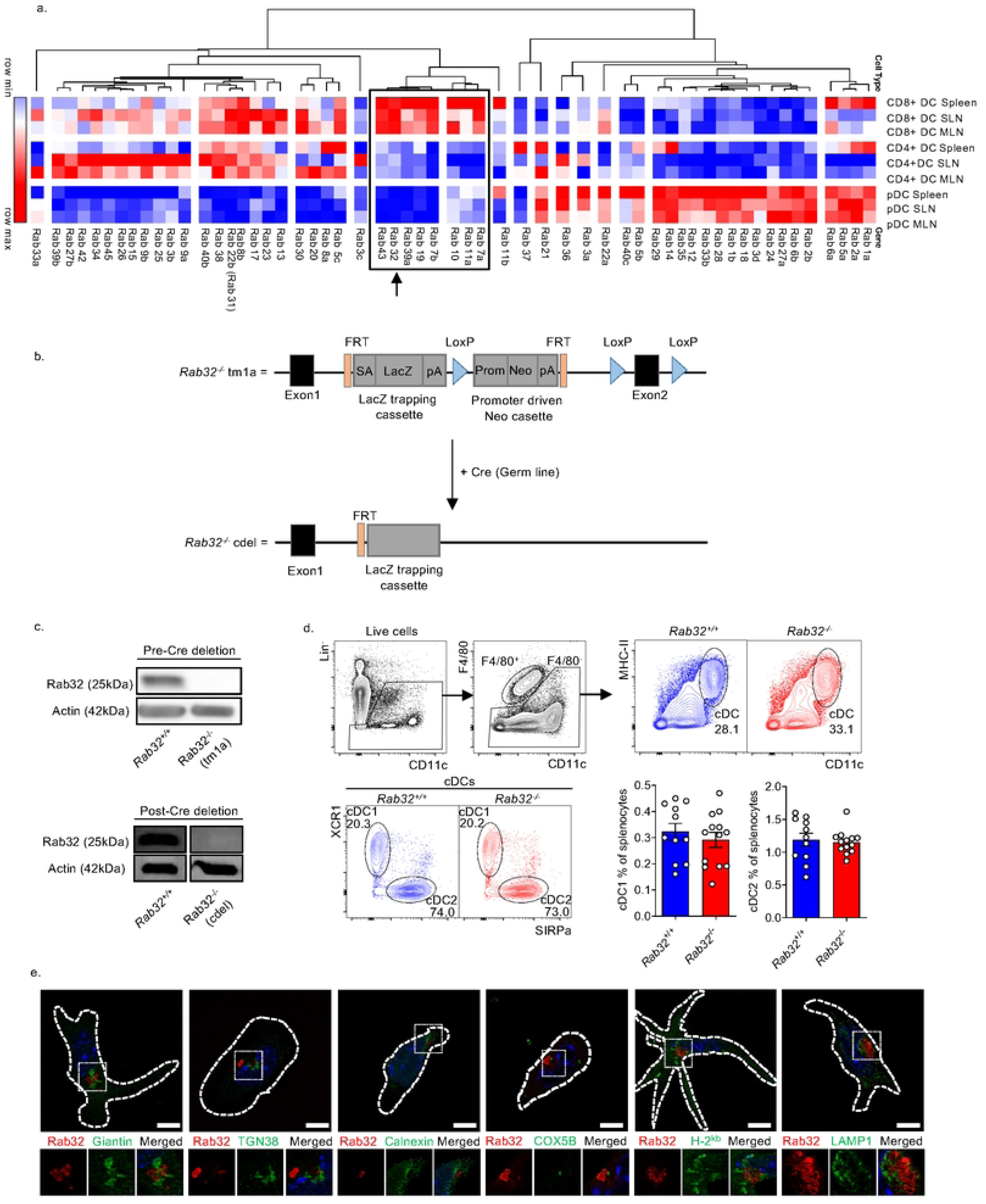
Rab32 is highly expressed in cDC1s but does not control their differentiation. a. Expression of Rab GTPases in various DC subsets in mice (microarray data from Immgen database). A group of Rabs – 7b, 19, 39a, 32 and 43 is differentially and highly expressed in cDC1s. b. The Rab32 tm1a have a promoter driven cassette present between the exon 1 and 2 or Rab32, which disrupts Rab32 protein expression. The tm1a mice were crossed with a germ line cre mice to create genetically deleted Rab32 KO mice (Rab32 cdel). c. Verification of the absence of Rab32 in the pre-Cre deletion Rab32 KO (tm1a) and post-Cre deletion Rab32 KO (cdel) mice by western blot. Analysis on tma1 and cdel performed on BMDCs and sorted splenic CD11c+XCR1+ cDC1s, respectively, with corresponding littermate control and actin loading control. d. FACS analysis of cDC populations in the spleens of naïve WT and KO mice. The Lin- consists of non-cDC markers including TER119, NK1.1, TCRb, GR-1 and CD19. Bar graphs depict the Quantification of XCR1+ cDC1s and SIRPa+ cDC2s populations. Data from 4 independent experiments. Each point represents an individual mouse. (WT n=11, KO n=13). Confocal microscopy projections of CD11c+ XCR1+ cDC1s sorted from the spleen of Rab32 WT mice, fixed and stained with antibodies against Rab32 along with various intracellular membrane markers- giatin (cis-Golgi), TGN38 (trans-Golgi), calnexin (Endoplasmic reticulum), COX5B (Mitochondria), H-2Kb and Lamp1 (Lysosomes). Scale bar - 7um.

To investigate the functional effects of Rab32 in a mice model, we used mice constitutively deficient in Rab32, named *Rab32^tm1a^*^/*tm1a*^. The Tm1a allele is based on a ‘Knock out first’ strategy, whereby the gene function is disrupted at the level of RNA processing. This is achieved by inserting a construct in the intron between exons 1 and 2 of Rab32. This construct consists of 2 cassettes – a ‘RNA trapping cassette’ containing LacZ reporter, and a floxed ‘promoter driven Neo cassette’ (Fig 1b). The trapping cassette contains a splice acceptor (SA) which traps the RNA transcript and fuses it with the LacZ reporter gene. It also contains a poly-adenylation (pA) sequence which truncates the RNA transcript, stopping the mRNA transcription downstream of the cassette. As anticipated, we show by western blot that Rab32 protein is not expressed in cells obtained from *Rab32^tm1a/tm1a^* mice (Fig 1c, upper blot). In addition, Rab32 exon 2 is floxed in the *Rab32^tm1a/tm1a^* mice, enabling its excision by cre-recombinase. We crossed the *Rab32^tm1a/tm1a^* mice with a germ-line ‘cre deleter’ mice to generate Rab32 cre-deleted (*Rab32^cdel/cdel^*) mice, in which exon 2 was ubiquitously deleted (fig 1b). As anticipated, *Rab32^cdel/cdel^* mice did not express Rab32 protein (Fig 1c, lower blot). Hence, both the *Rab32^tm1a/tm1a^* and the *Rab32^cdel/cdel^* mice were similarly deficient for Rab32 protein, and could serve as constitutive knockout models for further investigations on Rab32 function (termed herein as *Rab32*^-/-^ mice).

Using the *Rab32*^-/-^ mice, first we sought to determine if Rab32 affected the differentiation of cDC1s and cDC2s *in vivo*. To this end, we analyzed MHC-II^+^ CD11c^+^ XCR1^+^ cDC1s and SIRPa^+^ cDC2s from the spleens of *Rab32^+/+^* (littermate controls) or *Rab32^-/-^* mice by flow cytometry (Fig 1d). We found that the spleens of *Rab32^+/+^* or *Rab32^-/-^* mice had similar content in cDC1s and cDC2s. Moreover, Rab32 deficiency also did not affect the percentages of other myeloid and lymphoid cell populations in the spleen (Fig S1b). Hence, although Rab32 is highly expressed in cDC1s, it does not contribute to their differentiation pathway.

Next, we aimed to examine the intracellular localization of Rab32 in steady-state splenic cDC1s (Fig 1e). Unlike the cDC1-enriched Rab43, Rab32 did not colocalize with markers of cis- (Giantin) and trans-(TGN38) Golgi. Rab proteins have also been described to localize with the intracellular MHC-I storage compartments or lysosomal (LAMP1^+^) vesicles in DCs (15,23)(30). Rab32 was not found to colocalize with these compartments in primary splenic cDC1s. Previous studies have reported that Rab32 co-localises with the ER and mitochondria in cell lines (38). We did not observe a marked colocalization of Rab32 with the ER marker (calnexin) or Mitochondria (COX5B) in splenic cDC1s. By contrast, we found that Rab32 accumulated in perinuclear organelles of large size yet to be identified.

### Rab32 does not affect cross-presentation in short-term ex vivo assay

Having established that Rab32 does not control cDC development, we next wondered if Rab32 would control the overall ability of cDC1s to activate CD8^+^ T cells. From the phenotypic standpoint, cDCs from *Rab32^+/+^* or *Rab32^-/-^* mice did not display any detectable differences: MHC-I or MHC-II cell surface expression and the costimulatory markers CD40 and CD80 in naïve cDC1s were not affected by the deficiency of Rab32 in these cells (Fig 2a). We then examined whether Rab32 deficiency would affect the presentation of peptide epitope to CD8^+^ T cells. To this end, splenic cDC1s were sorted from *Rab32^+/+^* and *Rab32^-/-^* mice and co-cultured with CTV labeled OT-I cells (transgenic CD8^+^ T cells expressing a SIINFEKL specific TCR) in the presence of SIINFEKL peptide (normal OVA peptide presented on H-2K^b^ with high affinity to OT-I TCR) or SIIQFEKL peptide (mutated OVA peptide presented on H-2K^b^ with low affinity to OT-I TCR). Since OT-I cells are highly sensitive to SIINFEKL, minor differences in OT-I activation by *Rab32^+/+^* and *Rab32^-/-^* cDC1s could be better observed using SIIQFEKL. *Rab32^+/+^* and *Rab32^-/-^* cDC1s induced similar OT-I proliferation in response to the presentation of both these peptides, as measured by OT-I CTV divisions (Fig 2b). Moreover, OT-I activation, measured as IL-2 and IFNγ secretions in the culture supernatant, was also not affected by the deficiency of Rab32 in cDC1s (Fig 2c). Altogether we conclude that Rab32 is not essential in cDC1s for the presentation of peptide epitopes which do not require antigen processing in the cross presentation by MHC I pathway.

**Figure 2.**
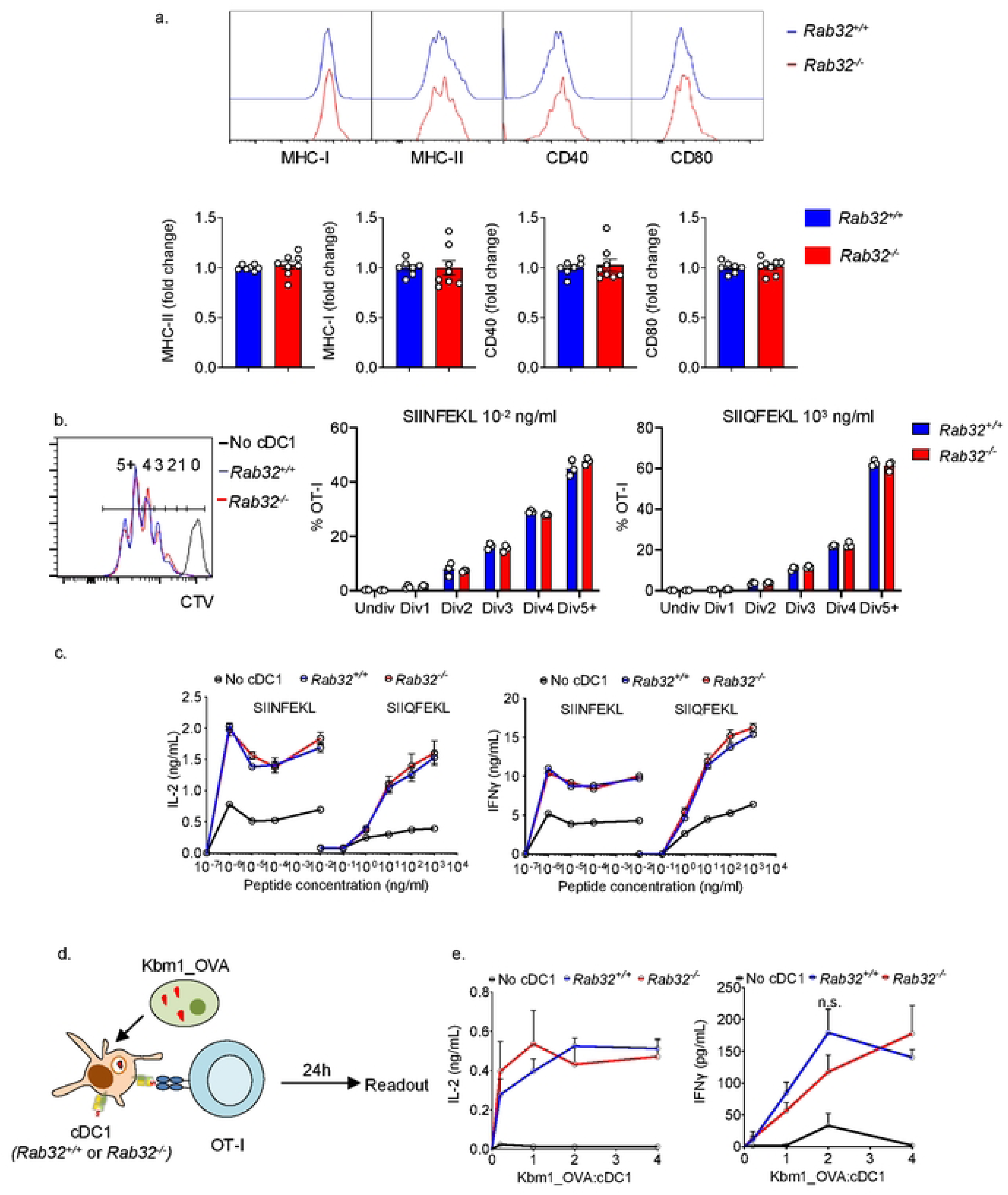
Rab32 does not control the cross-presentation of cell-associated antigens by cDC1s in vitro. a. FACS plots depict comparison MHC-I, MHC-II and activation marker (CD40 and CD80) expression on Rab32 WT and KO cDC1s. Bar graphs represent quantified data from multiple experiments. MHC-I, MHC-II and CD40 data from 2 independent experiments, CD80 data from 3 independent experiments. b. Spleen sorted CD11c+ XCR1+ cDC1s were co-cultured ex-vivo with CTV labelled and purified OT-I cells in the presence of different concentrations SIINFEKL (high affinity for OT-I TCR) or SIIQFEKL (mutated peptide, low affinity for OT-I TCR). FACS plot depicts OT-I divisions (CTV dilutions) in response to SIINFEKL peptide presentation by WT or KO cDC1s. OT-I divisions in response to SIINFEKL or SIIQFEKL presentation by WT or KO cDC1s were quantified. Data from 1 experiment with 3 individual cDC1 sortings from WT or KO mice each. c. In the same experiment, IL-2 (day1) and IFNγ (day3) secretions were quantified by ELISA. d. Experimental design for the ex-vivo cross-presentation assay. IL-2 and IFNγ secretions were measured as readout for cross-presentation. IL-2 and IFNγ secretions in presence of increasing Kbm1_OVA:cDC1 ration were quantified by ELISA. Data from 4 individual cDC1 sortings from WT or KO mice each.

Next, we examined if Rab32 would facilitate the process of antigen cross-presentation of cell-associated by splenic cDC1. *Rab32^+/+^* and *Rab32^-/-^* splenic cDC1s were co-cultured with OT-I cells *ex vivo*, in the presence of different numbers of K^bm1^_OVA cells (Fig 2d). IL-2 and IFNγ secretions by OT-I in the culture supernatant were measured as readouts for cross-presentation.

OT-I cells co-cultured with *Rab32^+/+^* or *Rab32^-/-^* cDC1s secreted similar quantities of both cytokines (Fig 2e). Based on these observations, we concluded that Rab32 expression in cDC1 is not essential for antigen cross-presentation by cDC1s *in vitro*.

### Rab32 controls the cross-priming against cell-associated antigens independently of its expression in CD8^+^ T cells

The lack of default in cross-presentation in *Rab32^-/-^* splenic cDC1s contradicted our initial hypothesis that cDC1-enriched Rab32 would the support the intracellular pathway for cross-presentation. We next wondered if Rab32 would control cDC1 cellular functions distinct from antigen processing, but required for efficient cross-priming *in vivo*.

To this end, we decided to test the response of adoptively transferred OT-I cells against K^bm1^_OVA challenge in *Rab32^+/+^* and *Rab32^-/-^* mice. CD45.1^+^/CD45.2^+^ OT-I cells were CTV labelled and injected into host CD45.2^+^ *Rab32^+/+^* or *Rab32^-/^* mice. 6-12h later, K^bm1^_OVA cells were *i.v.* injected in the same mice. 3 days later, the spleens of these mice were examined by flow cytometry to monitor OT-I activation and proliferation (Fig 3a). *Rab32^-/-^* mice showed consistently lower percentages of OT-I cells in the spleen for different numbers of K^bm1^_OVA cells injected (Fig 3b and c). Moreover, *Rab32^-/-^* mice had significantly lower percentages of effector (CTV^low^Cd62L^low^) OT-I cells compared to the *Rab32^+/+^* mice (Fig 3d and e). We conclude that Rab32 is essential for efficient cross-priming of CD8^+^ T cells against cell-associated antigens *in vivo*, by a process which is not T cell intrinsic.

**Figure 3.**
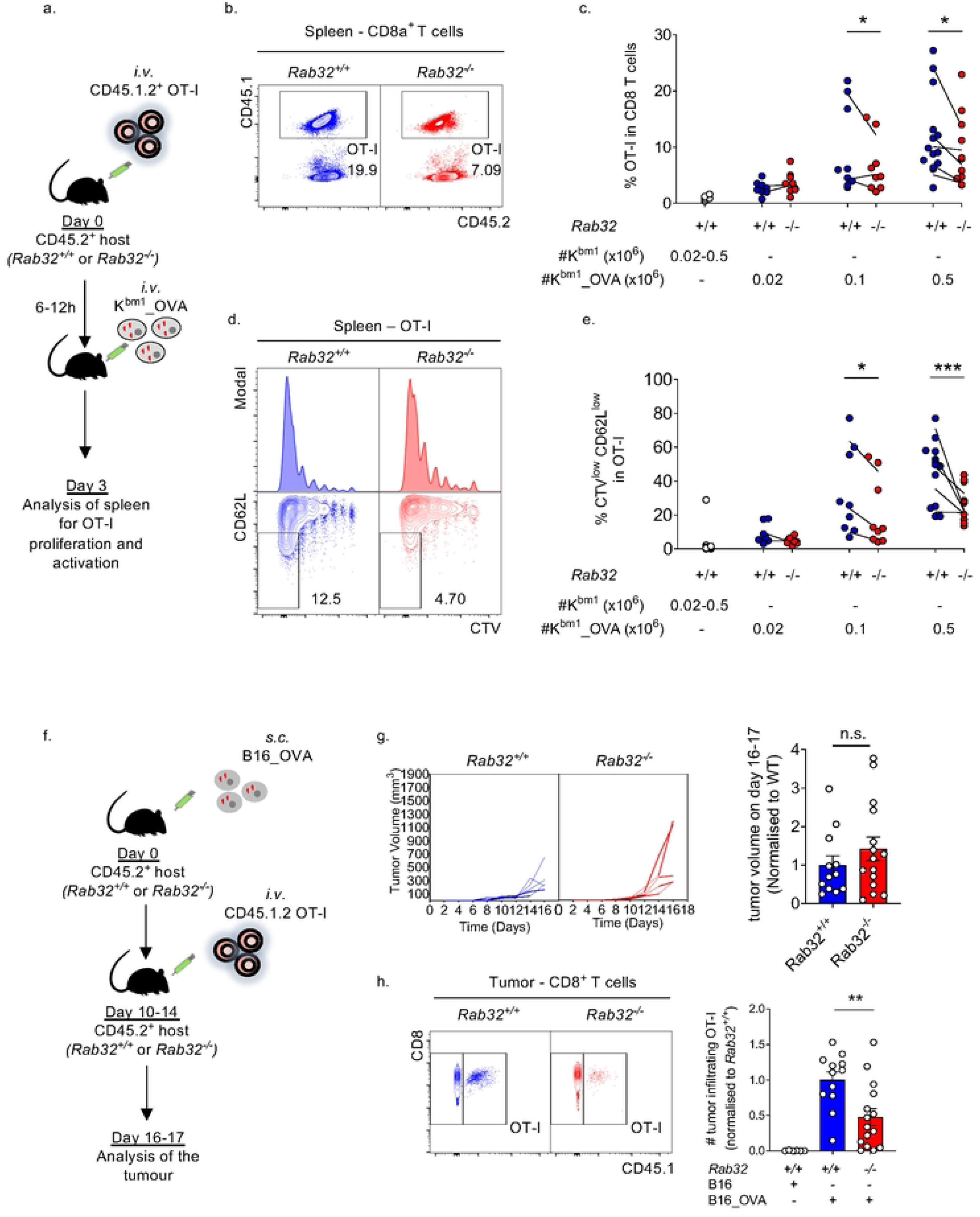
Rab32 controls the cross-priming against cell-associated antigens independently of its expression in CD8+ T cells. a. Experimental design for the OT-I adoptive transfer experiment. b. FACS panels depict a comparison of OT-I proliferation between WT and KO mice injected with 0.1x106 Kbm1_OVA. c. Quantification of the percentage of OT-I in the CD8+ T cell population in the spleens of WT and KO mice, for different numbers of injected Kbm1_OVA. Each point represents an individual mouse, and each line represents an individual experiment (*p<0.05, two-way ANOVA test). d. FACS panels depict a comparison of OT-I proliferation (CTV dilutions) and gain of effector function (loss of CD62L) between WT and KO mice. Quantification of the percentage of CTVlowCD62Llow cells in OT-I population in the spleens of WT and KO mice, for different numbers of injected Kbm1_OVA (*p<0.05, ***p<0.001, two-way ANOVA test).

In the next experiment, we wondered whether the role of Rab32 in the cross-priming of CD8+ T cells would apply to tumor antigens. B16_OVA melanomas were *s.c*. injected in CD45.2^+^ *Rab32^+/+^* and *Rab32^-/-^* mice. When the tumors grew to measurable sizes, CD45.1.2^+^ OT-I cells were *i.v.* injected in the same mice, 3-5 days later their pool was assessed in the tumor. (Fig 3f, gating strategy Fig S2a). *Rab32^-/-^* mice showed a tendency to grow larger tumors compared to *Rab32^+/+^* mice (Fig 3g). Moreover, *Rab32^-/-^* mice had a significantly lower number of OT-I cells in the tumors as compared to the *Rab32^+/+^* mice (Fig 3h). The *Rab32^-/-^* mice also had lower numbers of endogenous CD8^+^ T cells in the tumors, although the difference was not significant as compared to *Rab32^+/+^* mice (Fig S2b). Altogether, taking in account both endogenous and OT1, CD8+ T cell infiltration was significantly impaired in melanoma growing in Rab32-/- mice (Fig S2c). Hence, Rab32 significantly promoted the accrual of antigen-specific CD8^+^ T cells in B16 melanoma tumors.

Altogether, these observations lead us to conclude that Rab32 promotes the cross-priming against cell-associated antigens *in vivo* by virtue of its expression outside of the T cell compartment.

### Rab32 controls the Cross-priming against cell-associated antigens without affecting cDC1 maturation

The maturation status of cDCs is controls their ability to cross-prime effector cDC8^+^ T cells in vivo (43–45). Poly(I:C) is an agonist of both TLR-3 and MDA5 helicase and a strong inducer of cDC1 maturation (46)(47).Therefore, we sought to determine if Rab32 contributes to poly(I:C) induced cross-priming.

In the first line of experiments, we sought to test the maturation response of cDC1s to poly(I:C) *in vivo*. To this end, *Rab32^+/+^* and *Rab32^-/-^* mice were *i.p.* injected with 20ug poly(I:C), and 14h later, their spleens were examined by flow cytometry. *Rab32^+/+^* and *Rab32^-/-^* cDC1s similarly upregulated MHC-I, MHC-II as well as the co-stimulatory markers (CD40 and CD80) upon poly(I:C) stimulation (Fig 4a). In addition, blood serum from both *Rab32^+/+^* and *Rab32^-/-^* mice contained similar levels of the cytokine IL-12p40 (Fig S3a). Hence, we concluded that Rab32 does not affect the maturation or production of co-stimulatory signals (IL-12, *e.g*.) by cDC1s.

**Figure 4.**
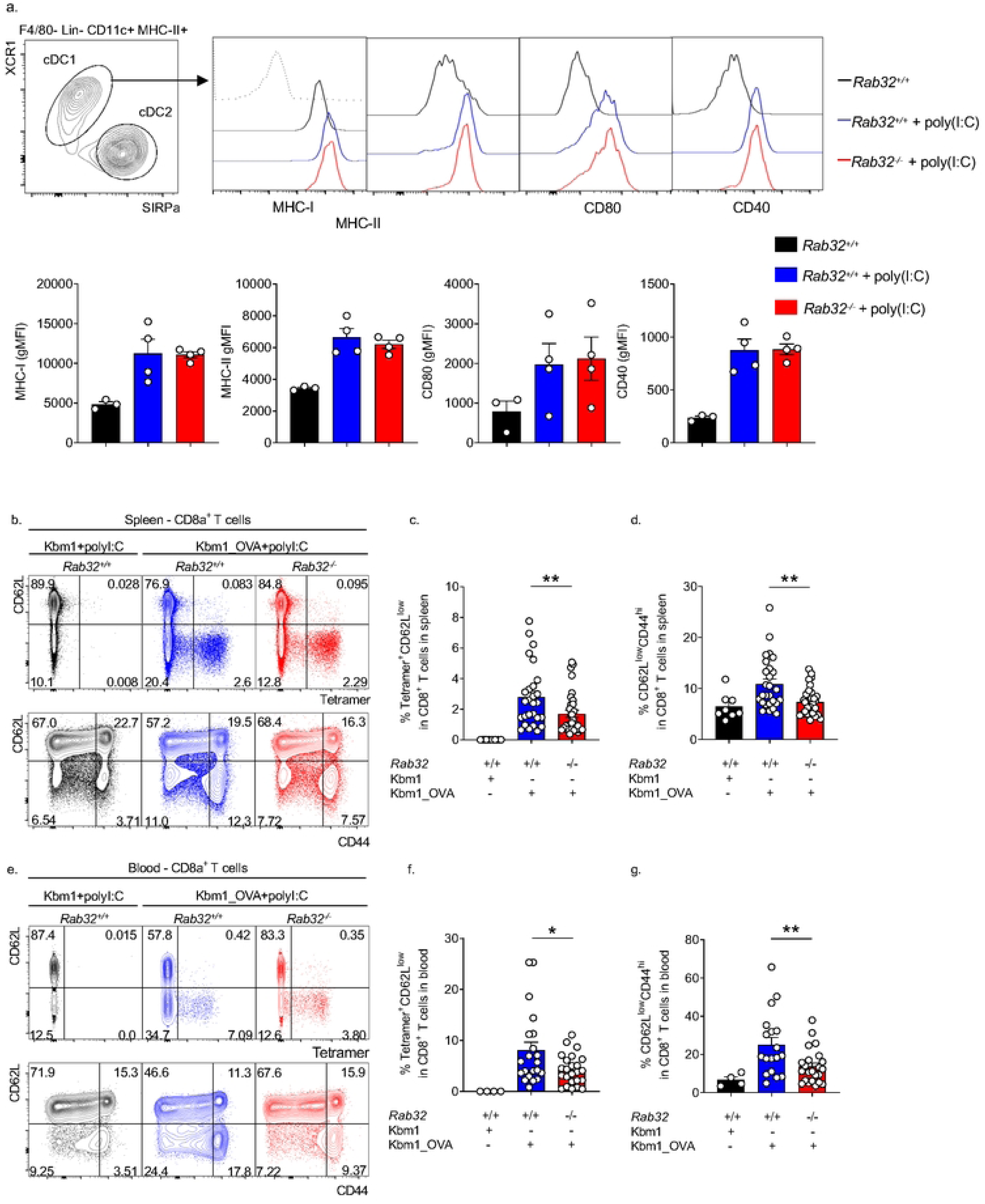
Rab32 is required for the efficient cross-priming of CD8+ T cells against cell- associated antigens in-vivo. a. Rab32 WT or KO mice were i.p. injected with 20ug polyI:C. 14h later, splenic cDC1s from the injected mice and non-injected WT controls were examined by FACS. The maturation phenotype of cDC1s was assessed by MHCI, MHCII, CD80 and CD40 levels. b. FACS plots depict the profile of endogenous CD8+ T cells in the spleens of Rab32 WT and KO mice, 10 days following immunisation with Kbm1_OVA+polyI:C. The upper panel depicts the OVA-specific (Tetramer+CD62Llow) CD8+ T cells, while lower panel depicts the effector (CD62LlowCD44hi) CD8+ T cells in the same spleen samples. c,d. Quantification of the percentages of Tetramer+ CD62Llow and CD44hi CD62Llow endogenous CD8+ T cells in the spleens of mice in each group. Data from 5 individual experiments. (Kbm1 control n=8, WT n=27, KO n=32, **p<0.01, Mann-Whitney test). e, f, g. Same as in b,c,d. but in the blood. Data from 3 individual experiments. (Kbm1 control n=4, WT n=12, KO n=18, *p<0.05, **p<0.01, unpaired t-test).

We next tested if Rab32 expression would be required for efficient poly(I:C)-dependent cross- priming. *Rab32^+/+^* and *Rab32^-/-^* mice were immunized with UV irradiated K^bm1^_OVA cells along with the TLR-3 agonist poly(I:C). Day 10 post-injection, the spleen and blood of these mice were examined for OVA-specific (Tetramer^+^CD62L^low^) and effector (CD62L^low^CD44^hi^) endogenous CD8^+^ T cells by flow cytometry (gating strategy – Fig S3b). *Rab32*^-/-^ mice showed significantly lower percentages of OVA-specific CD8^+^ T cells in the spleen, compared to *Rab32^+/+^* mice (Fig 4b upper panel and c). Moreover, *Rab32*^-/-^ mice also had lower percentages of effector CD8^+^ T cells in the spleen (Fig 4b lower panel and d). Similar observations were made in blood (Fig 4e-g, Fig. S3c). In both the genotypes, OVA-specific CD8^+^ T cells had similar effector status (Fig S3b).

Hence, we concluded that Rab32 is essential for efficient poly(I:C)-dependent cross-priming against cell-associated antigens *in vivo*.

## Discussion

In this report, we have analyzed the expression pattern of Rab GTPases in different resident cDC subsets. We report that Rab32 (together with Rab43, Rab39a, Rab7a, Rab7b, Rab11a, Rab10) is amongst the Rab GTPases highly expressed in resident cDC1 populations, as compared to cDC2s and pDCs (Fig 1a). We demonstrate that Rab32 is required for efficient activation of effector CD8^+^ T cells in response to the cross-presentation of cellular antigens *in vivo* (Fig XX). Interestingly, this default is only observed *in vivo*. This observation is reminiscent of the recently published report on the cDC1-specific Rab39a, which shows that Rab39a deficiency also does not affect cross-presentation by cDC1 *ex vivo*, yet it affects the cross-priming against cell-associated antigens *in vivo* (25). Our *ex vivo* experiments modeling CD8^+^ T cell cross-priming using MHC- incompetent donor cells did not evidence a default in the cross-presentation pathway of Rab32- deficient cDC1s purified from the spleen. This shows that Rab32 is not a key player in the intracellular pathways controlling the formation of peptide: MHC-I complexes after the phagocytosis of antigen. Such lack of function for Rab32, a well established lysosomal regulator might be explained by a compensatory mechanism. Indeed, certain Rab proteins are also known to be redundant in their functions. In the case of Rab32, a possible candidate is the closely related protein Rab38 (75% sequence similarity in mice). Rab32 and Rab38 are known to be essential for the biogenesis of melanosomes, which are lysosome-related organelles responsible for the synthesis and storage of melanin pigment in the body (48,49)(48)(50)(51). In the absence of either Rab38 or Rab32 activity, one can functionally compensate for the other, and hence, only a mild pigmentation phenotype is observed in single knockout cells (50). In Rab32 and Rab38 double knockout cells, the hypopigmentation phenotype is much more severe (51). The microarray data used in our report suggests that Rab38 is decently expressed in cDC1s (Fig S1a). Hence, such a functional redundancy may exist in cDC1s necessitating the analysis of cDC1s doubly deficient for Rab32 and Rab38.

Despite the lack of *- ex vivo* phenotype, *in vivo* experiments revealed a significant, despite partial default in the activation of CD8^+^ T cells specific for cell-associated antigens within *Rab32*^-/-^ mice. These results were found in 3 independent experimental settings involving distinct inflammatory signals: a) the cross-presentation of intravenously delivered K^bm1^_OVA cells, b) the cross-presentation of melanoma-associated model antigen (B16-OVA), c) the cross-presentation of intravenously delivered, necrotic fibroblasts adjuvanted in poly(I:C). In addition, we show that cDC maturation status - after poly(I:C) exposure is not controlled by Rab32 expression.

One might hypothesize that Rab32 expression would control the overall ability of cDC - to activate antigen-specific T cells (independently of antigen processing). However, our *ex vivo* assays did not reveal such default in Rab32-deficient cDC1s. Indeed, regardless of the antigen dose or affinity, TCR transgenic CD8^+^ T cells exposed to *Rab32^+/+^* or *Rab32*^-/-^ cDC1s in the presence of cognate peptide proliferated equally.

These data led to the hypothesis that Rab32 might control a biological process required for efficient T cell priming which is distinct from processing of cellular antigens and peptide: MHC-I complex formation. Such ‘processing-independent’ functions include, for instance, DC positioning within lymphoid organs (52), cytokine secretion (53), cDC1 migration (54), cDC1 survival during maturation (55). Future experiments should analyze the contribution of Rab32 to these processes essential for efficient cross-priming.

## Conclusion

In conclusion, our study reveals that the transcriptional landscape of cDC1 not only supports efficient antigen uptake and processing but also supports the implementation of other functions essential for efficient cross-priming but distinct from antigen processing and presentation by MHC-I. Altogether, this reveals novel roles for the Rab32 GTPase in the processes underlying the specialization of cDC1s in controlling CD8^+^ T cells responses against cell-associated antigens.

## Supplementary figures

Figure S1

a. Rab32 expression in multiples lineages. Rab32 is highly expressed in cDC1s, macrophages and neutrophils in comparison to T- and B-lymphocytes in mice (microarray data from Immgen database).

b. Percentages of various cell populations in the spleens of Rab32 WT and KO (tm1a) mice analysed by FACS. Data from 2 independent experiments (Mann-Whitney test).

Figure S2

a. Gating strategy for the analysis of tumour infiltrating OT-I cells. The Lin^-^ consists of non-cDC markers including TER119, NK1.1, TCRb, GR-1 and CD19.

b. And c. Quantification of the absolute numbers of endogenous CD8^+^ T cells and total CD8^+^ T cells (endogenous + OT-I) in the tumour in the same experiments as in the Figure 5c.

Figure S3

a. Serum IL12p40 in Rab32+/+ and Rab32-/- mice treated with polyI:C (one injection, 20ug, 14 hours earlier serum sampling).

b. Gating strategy for the analysis of endogenous, tetramer-OVA positive CD8+ T cells in mice immunized with Kbm1-OVA + poly(I:C) cells.

c. IFNg ELISPOT of blood leukocyte after SIINFEKL restimulation of Rab32^+/+^ and Rab32^-/-^ mice immunized with Kbm1-OVA + poly(I:C) cells.

## Acknowledgements

This work was supported by L’Agence Nationale de la Recherché (ANR) grant n. ANR-17-CE11- 0001-01 and ANR-18-IDEX-0001. PG is a CNRS investigator.

## Notes

### Competing Interest Statement

The authors have declared no competing interest.

## References

1. Bevan MJ. Minor H antigens introduced on H-2 different stimulating cells cross-react at the cytotoxic T cell level during in vivo priming. J Immunol. 1976 Dec;117(6):2233–8.

2. Bevan MJ. Cross-priming for a secondary cytotoxic response to minor H antigens with H-2 congenic cells which do not cross-react in the cytotoxic assay. J Exp Med. 1976 May 1;143(5):1283–8.

3. Bedoui S, Whitney PG, Waithman J, Eidsmo L, Wakim L, Caminschi I, et al. Cross-presentation of viral and self antigens by skin-derived CD103+ dendritic cells. Nat Immunol. 2009 May;10(5):488– 95.

4. Hildner K, Edelson BT, Purtha WE, Diamond M, Matsushita H, Kohyama M, et al. Batf3 deficiency reveals a critical role for CD8alpha+ dendritic cells in cytotoxic T cell immunity. Science. 2008 Nov 14;322(5904):1097–100.

5. Helft J, Manicassamy B, Guermonprez P, Hashimoto D, Silvin A, Agudo J, et al. Cross-presenting CD103+ dendritic cells are protected from influenza virus infection. J Clin Invest. 2012 Nov;122(11):4037–47.

6. Henri S, Poulin LF, Tamoutounour S, Ardouin L, Guilliams M, de Bovis B, et al. CD207+ CD103+ dermal dendritic cells cross-present keratinocyte-derived antigens irrespective of the presence of Langerhans cells. J Exp Med. 2010 Jan 18;207(1):189–206.

7. Schnorrer P, Behrens GM, Wilson NS, Pooley JL, Smith CM, El-Sukkari D, et al. The dominant role of CD8+ dendritic cells in cross-presentation is not dictated by antigen capture. Proc Natl Acad Sci U S A. 2006/06/30 ed. 2006 Jul 11;103(28):10729–34.

8. Poulin LF, Salio M, Griessinger E, Anjos-Afonso F, Craciun L, Chen JL, et al. Characterization of human DNGR-1 ^+^ BDCA3 ^+^ leukocytes as putative equivalents of mouse CD8α ^+^ dendritic cells. The Journal of Experimental Medicine. 2010 Jun 7;207(6):1261–71.

9. Jongbloed SL, Kassianos AJ, McDonald KJ, Clark GJ, Ju X, Angel CE, et al. Human CD141+ (BDCA-3)+ dendritic cells (DCs) represent a unique myeloid DC subset that cross-presents necrotic cell antigens. Journal of Experimental Medicine. 2010 Jun 7;207(6):1247–60.

10. Balan S, Ollion V, Colletti N, Chelbi R, Montanana-Sanchis F, Liu H, et al. Human XCR1+ Dendritic Cells Derived In Vitro from CD34+ Progenitors Closely Resemble Blood Dendritic Cells, Including Their Adjuvant Responsiveness, Contrary to Monocyte-Derived Dendritic Cells. The Journal of Immunology. 2014 Aug 15;193(4):1622–35.

11. Bachem A, Güttler S, Hartung E, Ebstein F, Schaefer M, Tannert A, et al. Superior antigen cross- presentation and XCR1 expression define human CD11c+CD141+ cells as homologues of mouse CD8+ dendritic cells. Journal of Experimental Medicine. 2010 Jun 7;207(6):1273–81.

12. den Haan JMM, Bevan MJ. Constitutive versus Activation-dependent Cross-Presentation of Immune Complexes by CD8 ^+^ and CD8 ^−^ Dendritic Cells In Vivo. The Journal of Experimental Medicine. 2002 Sep 16;196(6):817–27.

13. Desch AN, Randolph GJ, Murphy K, Gautier EL, Kedl RM, Lahoud MH, et al. CD103+ pulmonary dendritic cells preferentially acquire and present apoptotic cell-associated antigen. J Exp Med. 2011 Aug 29;208(9):1789–97.

14. Cebrian I, Visentin G, Blanchard N, Jouve M, Bobard A, Moita C, et al. Sec22b regulates phagosomal maturation and antigen crosspresentation by dendritic cells. Cell. 2011 Dec 9;147(6):1355–68.

15. Nair-Gupta P, Baccarini A, Tung N, Seyffer F, Florey O, Huang Y, et al. TLR signals induce phagosomal MHC-I delivery from the endosomal recycling compartment to allow cross-presentation. Cell. 2014 Jul 31;158(3):506–21.

16. Savina A, Jancic C, Hugues S, Guermonprez P, Vargas P, Moura IC, et al. NOX2 controls phagosomal pH to regulate antigen processing during crosspresentation by dendritic cells. Cell. 2006 Jul 14;126(1):205–18.

17. Zehner M, Marschall AL, Bos E, Schloetel JG, Kreer C, Fehrenschild D, et al. The translocon protein Sec61 mediates antigen transport from endosomes in the cytosol for cross-presentation to CD8(+) T cells. Immunity. 2015 May 19;42(5):850–63.

18. Helft J, Böttcher J, Chakravarty P, Zelenay S, Huotari J, Schraml BU, et al. GM-CSF Mouse Bone Marrow Cultures Comprise a Heterogeneous Population of CD11c+MHCII+ Macrophages and Dendritic Cells. Immunity. 2015 Jun 16;42(6):1197–211.

19. Briseño CG, Haldar M, Kretzer NM, Wu X, Theisen DJ, Kc W, et al. Distinct Transcriptional Programs Control Cross-Priming in Classical and Monocyte-Derived Dendritic Cells. Cell Rep. 2016 Jun 14;15(11):2462–74.

20. Sancho D, Joffre OP, Keller AM, Rogers NC, Martínez D, Hernanz-Falcón P, et al. Identification of a dendritic cell receptor that couples sensing of necrosis to immunity. Nature. 2009 Apr 16;458(7240):899–903.

21. Canton J, Blees H, Henry CM, Buck MD, Schulz O, Rogers NC, et al. The receptor DNGR-1 signals for phagosomal rupture to promote cross-presentation of dead-cell-associated antigens. Nat Immunol. 2021 Feb;22(2):140–53.

22. Bougnères L, Helft J, Tiwari S, Vargas P, Chang BHJ, Chan L, et al. A Role for Lipid Bodies in the Cross-presentation of Phagocytosed Antigens by MHC Class I in Dendritic Cells. Immunity. 2009 Aug;31(2):232–44.

23. Kretzer NM, Theisen DJ, Tussiwand R, Briseño CG, Grajales-Reyes GE, Wu X, et al. RAB43 facilitates cross-presentation of cell-associated antigens by CD8α+ dendritic cells. J Exp Med. 2016 Dec 12;213(13):2871–83.

24. Theisen DJ, Davidson JT, Briseño CG, Gargaro M, Lauron EJ, Wang Q, et al. WDFY4 is required for cross-presentation in response to viral and tumor antigens. Science. 2018 09;362(6415):694–9.

25. Cruz FM, Colbert JD, Rock KL. The GTPase Rab39a promotes phagosome maturation into MHC-I antigen-presenting compartments. EMBO J. 2020 Jan 15;39(2):e102020.

26. Rodríguez-Silvestre P, Laub M, Krawczyk PA, Davies AK, Schessner JP, Parveen R, et al. Perforin-2 is a pore-forming effector of endocytic escape in cross-presenting dendritic cells. Science. 2023 Jun 23;380(6651):1258–65.

27. Gonzales GA, Huang S, Wilkinson L, Nguyen JA, Sikdar S, Piot C, et al. The pore-forming apolipoprotein APOL7C drives phagosomal rupture and antigen cross-presentation by dendritic cells. Sci Immunol. 2024 Nov;9(101):eadn2168.

28. Lin ML, Zhan Y, Proietto AI, Prato S, Wu L, Heath WR, et al. Selective suicide of cross-presenting CD8+ dendritic cells by cytochrome c injection shows functional heterogeneity within this subset. Proc Natl Acad Sci U S A. 2008 Feb 26;105(8):3029–34.

29. Dudziak D, Kamphorst AO, Heidkamp GF, Buchholz VR, Trumpfheller C, Yamazaki S, et al. Differential antigen processing by dendritic cell subsets in vivo. Science. 2007/01/06 ed. 2007 Jan 5;315(5808):107–11.

30. Zou L, Zhou J, Zhang J, Li J, Liu N, Chai L, et al. The GTPase Rab3b/3c-positive recycling vesicles are involved in cross-presentation in dendritic cells. Proc Natl Acad Sci U S A. 2009 Sep 15;106(37):15801–6.

31. Savina A, Peres A, Cebrian I, Carmo N, Moita C, Hacohen N, et al. The small GTPase Rac2 controls phagosomal alkalinization and antigen crosspresentation selectively in CD8(+) dendritic cells. Immunity. 2009 Apr 17;30(4):544–55.

32. Li Y, Wang Y, Zou L, Tang X, Yang Y, Ma L, et al. Analysis of the Rab GTPase Interactome in Dendritic Cells Reveals Anti-microbial Functions of the Rab32 Complex in Bacterial Containment. Immunity. 2016 Feb 16;44(2):422–37.

33. Spanò S, Galán JE. A Rab32-dependent pathway contributes to Salmonella typhi host restriction. Science. 2012 Nov 16;338(6109):960–3.

34. Spanò S, Gao X, Hannemann S, Lara-Tejero M, Galán JE. A Bacterial Pathogen Targets a Host Rab- Family GTPase Defense Pathway with a GAP. Cell Host Microbe. 2016 Feb 10;19(2):216–26.

35. Solano-Collado V, Rofe A, Spanò S. Rab32 restriction of intracellular bacterial pathogens. Small GTPases. 2018 May 4;9(3):216–23.

36. Chen M, Sun H, Boot M, Shao L, Chang SJ, Wang W, et al. Itaconate is an effector of a Rab GTPase cell-autonomous host defense pathway against Salmonella. Science. 2020 Jul 24;369(6502):450–5.

37. Xie X, Ni Q, Zhou D, Wan Y. Rab32-related antimicrobial pathway is involved in the progression of dextran sodium sulfate-induced colitis. FEBS Open Bio. 2018 Oct;8(10):1658–68.

38. Bui M, Gilady SY, Fitzsimmons REB, Benson MD, Lynes EM, Gesson K, et al. Rab32 modulates apoptosis onset and mitochondria-associated membrane (MAM) properties. J Biol Chem. 2010 Oct 8;285(41):31590–602.

39. Ortiz-Sandoval CG, Hughes SC, Dacks JB, Simmen T. Interaction with the effector dynamin-related protein 1 (Drp1) is an ancient function of Rab32 subfamily proteins. Cell Logist. 2014;4(4):e986399.

40. Alto NM, Soderling J, Scott JD. Rab32 is an A-kinase anchoring protein and participates in mitochondrial dynamics. J Cell Biol. 2002 Aug 19;158(4):659–68.

41. Gerondopoulos A, Langemeyer L, Liang JR, Linford A, Barr FA. BLOC-3 mutated in Hermansky- Pudlak syndrome is a Rab32/38 guanine nucleotide exchange factor. Curr Biol. 2012 Nov 20;22(22):2135–9.

42. Ohishi Y, Kinoshita R, Marubashi S, Ishida M, Fukuda M. The BLOC-3 subunit HPS4 is required for activation of Rab32/38 GTPases in melanogenesis, but its Rab9 activity is dispensable for melanogenesis. J Biol Chem. 2019 Apr 26;294(17):6912–22.

43. Bonifaz L, Bonnyay D, Mahnke K, Rivera M, Nussenzweig MC, Steinman RM. Efficient Targeting of Protein Antigen to the Dendritic Cell Receptor DEC-205 in the Steady State Leads to Antigen Presentation on Major Histocompatibility Complex Class I Products and Peripheral CD8+ T Cell Tolerance. Journal of Experimental Medicine. 2002 Dec 16;196(12):1627–38.

44. Hawiger D, Inaba K, Dorsett Y, Guo M, Mahnke K, Rivera M, et al. Dendritic Cells Induce Peripheral T Cell Unresponsiveness under Steady State Conditions in Vivo. Journal of Experimental Medicine. 2001 Sep 17;194(6):769–80.

45. Steinman RM, Hawiger D, Nussenzweig MC. Tolerogenic dendritic cells. Annu Rev Immunol. 2003;21:685–711.

46. Pantel A, Teixeira A, Haddad E, Wood EG, Steinman RM, Longhi MP. Direct type I IFN but not MDA5/TLR3 activation of dendritic cells is required for maturation and metabolic shift to glycolysis after poly IC stimulation. PLoS Biol. 2014 Jan;12(1):e1001759.

47. Schulz O, Diebold SS, Chen M, Näslund TI, Nolte MA, Alexopoulou L, et al. Toll-like receptor 3 promotes cross-priming to virus-infected cells. Nature. 2005 Feb 24;433(7028):887–92.

48. Bultema JJ, Boyle JA, Malenke PB, Martin FE, Dell’Angelica EC, Cheney RE, et al. Myosin vc interacts with Rab32 and Rab38 proteins and works in the biogenesis and secretion of melanosomes. J Biol Chem. 2014 Nov 28;289(48):33513–28.

49. Bultema JJ, Ambrosio AL, Burek CL, Di Pietro SM. BLOC-2, AP-3, and AP-1 proteins function in concert with Rab38 and Rab32 proteins to mediate protein trafficking to lysosome-related organelles. J Biol Chem. 2012 Jun 1;287(23):19550–63.

50. Loftus SK, Larson DM, Baxter LL, Antonellis A, Chen Y, Wu X, et al. Mutation of melanosome protein RAB38 in chocolate mice. Proc Natl Acad Sci U S A. 2002 Apr 2;99(7):4471–6.

51. Wasmeier C, Romao M, Plowright L, Bennett DC, Raposo G, Seabra MC. Rab38 and Rab32 control post-Golgi trafficking of melanogenic enzymes. J Cell Biol. 2006 Oct 23;175(2):271–81.

52. Eickhoff S, Brewitz A, Gerner MY, Klauschen F, Komander K, Hemmi H, et al. Robust Anti-viral Immunity Requires Multiple Distinct T Cell-Dendritic Cell Interactions. Cell. 2015 Sep 10;162(6):1322–37.

53. Chiaruttini G, Piperno GM, Jouve M, De Nardi F, Larghi P, Peden AA, et al. The SNARE VAMP7 Regulates Exocytic Trafficking of Interleukin-12 in Dendritic Cells. Cell Rep. 2016 Mar 22;14(11):2624–36.

54. Bretou M, Sáez PJ, Sanséau D, Maurin M, Lankar D, Chabaud M, et al. Lysosome signaling controls the migration of dendritic cells. Sci Immunol. 2017 Oct 27;2(16):eaak9573.

55. Fuertes Marraco SA, Scott CL, Bouillet P, Ives A, Masina S, Vremec D, et al. Type I interferon drives dendritic cell apoptosis via multiple BH3-only proteins following activation by PolyIC in vivo. PLoS One. 2011;6(6):e20189.

